# Deviation of Dyar’s rule in the post-embryonic development in millipedes – a comprehensive analysis

**DOI:** 10.1101/2020.08.23.263848

**Authors:** Somnath Bhakat

## Abstract

Dyar’s value on the basis of length and width of nine polydesmid and 15 non-polydesmid millipede species were calculated. The value of polydesmid millipede ranged from 1.50 to 1.78 and that of non-polydesmid millipede ranged from 1.08 to 1.45. Weight progression factor was determined for nine species of millipede (two polydesmid and seven non-polydesmids). The result showed that the mean weight progression factor in polydesmid is 2.54 while that of non-polydesmid is 1.95. Both the results showed that Dyar’s value in polydesmida is significantly higher compared to that of non-polydesmida. In polydesmida, the number of stadium is only eight with higher Dyar’s value (mean 1.61) while in non-polydesmida, where number of stadium is more than eight have lower Dyar’s value (mean 1.23). As in other arthropods, Dyar’s value is inversely proportional to the number of stadium in millipede.

The present study also affirmed Enders’ hypothesis in favour of adaptive importance of Dyar’s rule. Deviations from Dyar’s constant in these two group also support Crossby’s growth rule. The variation of Dyar’s value in these two groups of millipede is related to the development time and habitat utilization. The variation of weight progression factor in these two groups is also linked to the development time as observed in other arthropods.

## Introduction

Arthropods are covered by a cuticle that mainly functions as an exoskeleton. In most arthropods, these sclerotized exoskeletal structures grow in a successive manner at each moult. The growth of the sclerotized parts can be estimated by calculating their size in successive instars. From a study of the larvae of some 28 species of Lepidoptera, Dyar (1890) concluded that size increase of hard structure in caterpillar follows a geometric progression. The so called Dyar’s rule has been confirmed for a lot of arthropods (Taylor, 1931; Elliott, 2009) and it has occasionally been used to determine the number of instars of some species (Gaines and Campbell, 1935). Several authors, however, have reported deviations from Dyar’s rule (Drooz, 1965; Hoxie and Wellso, 1974; Jobin et al, 1992; Andersson, 1976, 1978b, 1980; Albert, 1982, Cole, 1980).

Dyar (1890) also found that the factor by which caterpillars grow in successive moult is the same for all species, about 1.26 when body length (BL) and about 2 when weight is considered. Comparison of Dyar’s constant should be made only on sclerotized parts because these do not change within an instar. For the sake of comparison these measurements should always be related to BL and Dyar’s value calculated for BL. The total body length is a measurement that is frequently used since it has an obvious biological significance (Buxton, 1938; Clark and Harsh, 1939; Matsuda, 1963). Observations on the increase of body length in successive stadia in millipede indicate that the rule used in Lepidoptera larva can also be used as a unit for measuring growth in this arthropod group.

In this paper it will be tested by an analysis of literature data and own values for two non-polydesmid (Spirobolid and Spirostreptid) millipede whether Dyar’ rule really works for millipedes. In particular, in this study, following hypotheses have been tested:

1. That if the millipede obeys the Dyar’s constant during post-embryonic development.
2. That if there is significant variation in Dyar’ value in the millipedes having short and long developmental time (polydesmid vs. non-polydesmid).
3. That if there is significant difference in weight progression factor of male and female.
4. That if Dyar’s value indicates any habitat segregation and life history strategy in these two groups.

## Material and Methods

In the literature most of the authors separated different stadia of millipedes by number of podous and apodous segments, leg pairs or ocelli present in each row except a few who mention the size (length and width) of each stadium. Lengths and widths of different stadia of *G. malayus* and *T. lumbricinus* were determined as described by Blower and Gabbut (1964), Blower and Miller (1977) and Bhakat (1987). Stadia were confirmed by counting the rows of ocelli followed by Vachon (1947). Weight of each stadium was measured by digital single pan balance with an accuracy of 0.1 mg. Different stadia of these two species were collected from different fields in and around Suri (87°32’00’’E, 23°55’00’’N) (District-Birbhum, West Bengal, India). Length, width and weight of these two species are listed in Table 1. Dyar’s constant or efficiency (r) can be calculated from the following formula:

**Table 1.** Characteristics of different stadia of *G. malayus* and *T. lumbricinus* (M-male, F-female).

| Species | *G. malayus* |  |  | *T. lumbricinus* |  |  |
| --- | --- | --- | --- | --- | --- | --- |
| Stadia | Length cm | Width mm | Stadia | Length mm | Width mm | Weight mg |
| IV | 5.8 | 3.9 | II | 3.2 – 5.9 |  |  |
| V | 6.4 | 4.2 | III | 5.7 – 6.8 |  | 1.19 – 2.15 |
| VI | 7.1 | 4.5 | IV | 6.3 – 7.5 | 0.90 – 1.10 | 6.45 – 10.25 |
| VII | 7.8 | 4.8 | V M | 8.6 – 9.2 | 1.12 – 1.14 | 14.05 – 20.85 |
| VIII | 8.3 | 5.3 | F | 9.0 – 9.5 | 1.15 – 1.50 | 14.50 – 21.50 |
| IX | 9.0 | 5.9 | VI M | 14.5 – 22.0 | 1.51 – 2.15 | 38.74 – 91.35 |
| X | 9.2 | 6.2 | F | 16.5 – 23.5 | 1.95 – 2.60 | 56.74 – 105.30 |
| XI | 10.1 | 6.4 | VII M | 24.5 – 30.0 | 2.25 – 2.90 | 98.35 – 183.74 |
| XII | 10.8 | 7.0 | F | 30.0 – 32.5 | 2.65 – 3.40 | 136.09 – 229.50 |
| XIII | 11.2 | 7.5 | VIII M | 30.2 – 36.0 | 2.60 – 3.15 | 216.35 – 350.50 |
| XIV | 11.8 | 8.2 | F | 31.5 – 38.0 | 3.00 – 3.70 | 225.55 – 595.74 |
| XV | 13.2 | 9.0 | IX M | 42.5 – 51.5 | 3.55 – 3.85 | 487.35 – 810.30 |
|  |  |  | F | 45.5 – 52.0 | 3.60 – 4.10 | 630.35 – 1009.70 |

r = X_i+1_/X_i_ Where, X_i_ is the value of a linear size variable at the ith stadium (body length or width) and X_i+1_ is the value of the same variable at the following stadia.

In the present study, Dyar’s rule is applied on 10 species of Polydesmida and 18 species of non-Polydesmida (Table 2).

**Table 2.**
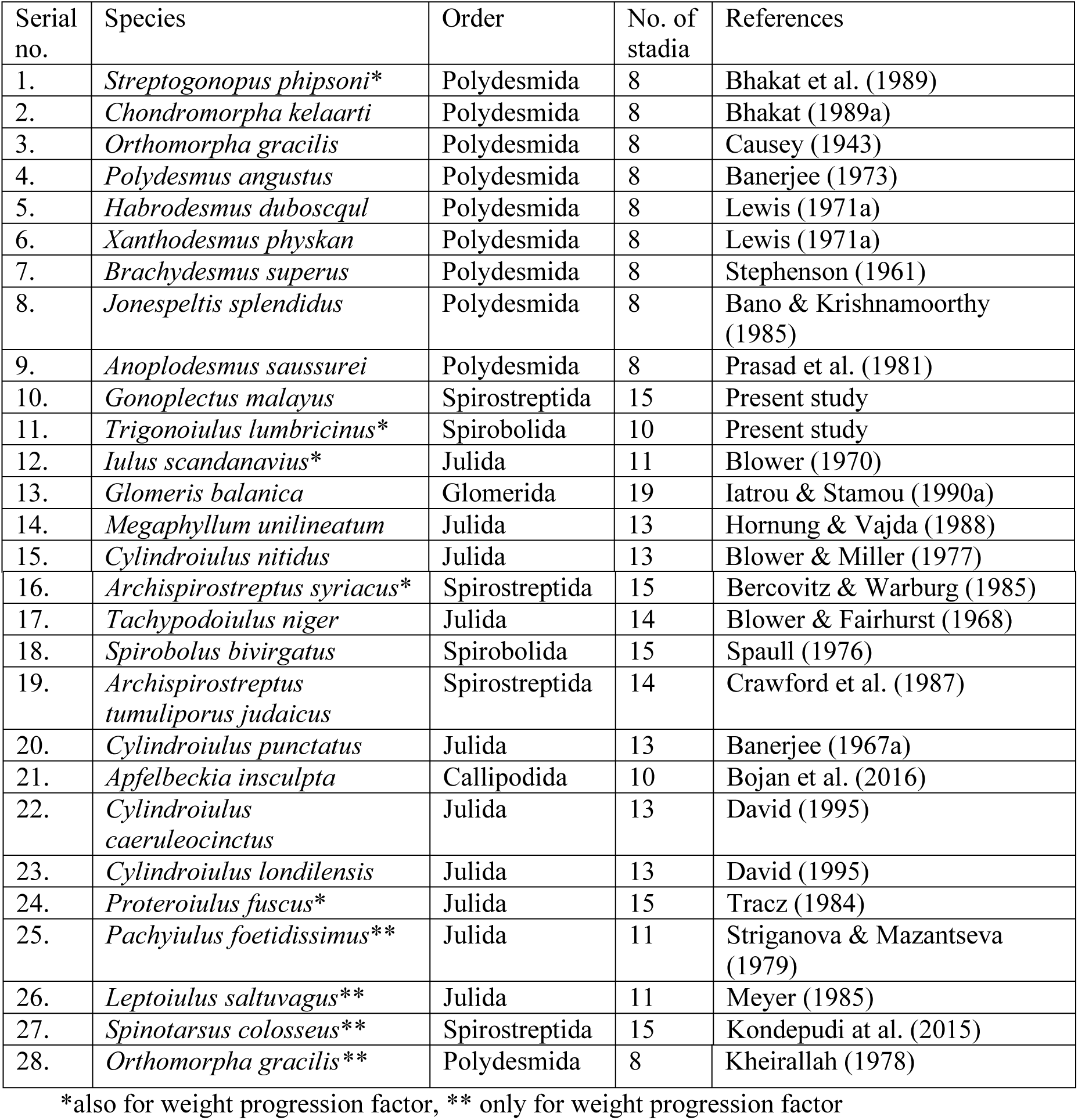
List of millipede species used for calculation of Dyar’s constant.

Weight progression factor on the basis of body weight could only be determined for eight species of millipedes by the above mentioned formula (where, X_i_ = weight of individual stadium).

## Results

Dyar’s values for polydesmid and non-polydesmid millipede are presented in Table 3 and 4 respectively. Dyar’s value of all the species of polydesmida is >1.5 with a mean value 1.61 (Table 3) but that of non-polydesmid millipede is <1.5 with a mean value 1.23 (Table 4). Dyar’s value for polydesmid is significantly higher than a postulated value of 1.26 (Przibram and Megusar, 1912; and Bodenheimer, 1927) or 1.28 (Horn and May, 1977 and Maiorana, 1978) though in case of non-polydesmid millipede it is nearer to the postulated value. The difference of Dyar’s value in those two groups of millipede is highly significant (P < 0.001).

**Table 3.** Dyar’s value of different stadia in different species of Polydesmid millipede (L = length, W = width).

| Species/stadia |  | II | III | IV | V | VI | VII | VIII | Mean |
| --- | --- | --- | --- | --- | --- | --- | --- | --- | --- |
| *S. phipsoni* | L | 1.34 | 1.48 | 2.11 | 1.35 | 1.42 | 1.66 | 1.99 | 1.62 |
|  | W | 1.69 | 1.55 | 1.29 | 1.27 | 1.82 | 1.77 | 1.50 | 1.56 |
| *C. kelaarti* | L | - | 3.24 | 1.18 | 1.88 | 1.12 | 1.55 | 1.11 | 1.68 |
|  | W | - | 1.92 | 1.29 | 1.79 | 1.19 | 1.24 | 1.88 | 1.55 |
| *O. gracilis* | L | 2.65 | 2.67 | 1.22 | 1.20 | 1.46 | 1.69 | 1.56 | 1.78 |
| *P. angustus* | L | 1.62 | 1.81 | 1.50 | 1.18 | 1.90 | 1.40 | - | 1.57 |
| *H. duboscqui* | W | - | 1.46 | 1.59 | 1.67 | 1.60 | 1.45 | 1.63 | 1.57 |
| *X. physkans* | W | - | 1.50 | 1.57 | 1.49 | 1.44 | 1.71 | 1.65 | 1.56 |
| *B. superus* | L | 1.75 | 1.57 | 1.50 | 1.39 | 1.28 | - | - | 1.50 |
| *J. splendidus* | L | 2.07 | 1.62 | 1.46 | 1.56 | 1.96 | 1.28 | 1.60 | 1.65 |
| *A. saussurei* | L | 2.45 | 1.59 | 1.66 | 1.62 | 1.36 | 1.65 | 2.07 | 1.77 |
|  | W | 1.18 | 1.30 | 1.27 | 1.65 | 1.31 | 1.47 | 2.48 | 1.52 |

**Table 4.** Dyar’s value of different stadia in different species of non-Polydesmid millipede (L = length, W = width)

| Species | *G. malayus* |  | *T. lumbricinus* |  | *G. balanica* |  | *M. unilineatum* | *C. nitidus* |  |
| --- | --- | --- | --- | --- | --- | --- | --- | --- | --- |
| Stadia | L | W | L | W | L | W | W | L | W |
| II |  |  |  |  | 1.16 |  |  |  |  |
| III |  |  | 1.47 |  | 1.23 |  |  |  |  |
| IV |  |  | 1.11 |  | 1.33 |  |  |  |  |
| V | 1.10 | 1.08 | 1.33 | 1.23 | 1.23 |  | 1.20 |  |  |
| VI | 1.11 | 1.07 | 2.10 | 1.66 |  | 1.16 | 1.24 |  |  |
| VII | 1.10 | 1.07 | 1.57 | 1.38 |  | 1.15 | 1.17 | 1.23 | 1.16 |
| VIII | 1.06 | 1.10 | 1.16 | 1.12 |  | 1.14 | 1.63 | 1.19 | 1.18 |
| IX | 1.08 | 1.11 | 1.42 | 1.22 |  | 1.11 |  | 1.18 | 1.12 |
| X | 1.02 | 1.05 |  |  |  | 1.06 |  | 1.14 | 1.12 |
| XI | 1.10 | 1.03 |  |  |  | 1.08 |  | 1.15 | 1.08 |
| XII | 1.07 | 1.09 |  |  |  | 1.06 |  | 1.14 | 1.00 |
| XIII | 1.04 | 1.07 |  |  |  | 1.07 |  | 1.06 | 1.13 |
| XIV | 1.05 | 1.09 |  |  |  | 1.06 |  |  |  |
| XV | 1.12 | 1.10 |  |  |  | 1.06 |  |  |  |
| XVI |  |  |  |  |  | 1.05 |  |  |  |
| XVII |  |  |  |  |  | 1.05 |  |  |  |
| XVIII |  |  |  |  |  | 1.04 |  |  |  |
| XIX |  |  |  |  |  | 1.03 |  |  |  |
| Mean | 1.08 | 1.08 | 1.45 | 1.32 | 1.24 | 1.08 | 1.31 | 1.15 | 1.11 |

Table 4. Contd.
| Species | *I. scandinavicus* |  | *A. syriacus* |  | *T. niger* | *S. bivirgatus* | *A. tumuliporus judaicus* |
| --- | --- | --- | --- | --- | --- | --- | --- |
| Stadia | L | W | L | W | L | L | W |
| II |  |  |  |  |  | 1.53 |  |
| III |  |  | 1.20 | 1.00 |  | 1.69 |  |
| IV |  | 1.20 | 1.25 | 1.21 |  | 1.42 |  |
| V | 2.13 | 1.20 | 1.27 | 1.21 |  | 1.48 |  |
| VI | 1.22 | 1.19 | 1.37 | 1.20 |  | 1.13 |  |
| VII | 1.32 | 1.27 |  |  |  | 1.35 |  |
| VIII | 1.32 | 1.18 |  |  |  | 1.39 | 1.20 |
| IX | 1.26 | 1.22 |  |  | 1.19 | 1.11 | 1.06 |
| X | 1.30 | 1.22 |  |  | 1.18 | 1.06 | 1.18 |
| XI | 1.10 | 1.11 |  |  | 1.14 | 1.05 | 1.13 |
| XII |  |  |  |  | 1.08 | 1.09 | 1.03 |
| XIII |  |  |  |  |  | 1.13 | 1.01 |
| XIV |  |  |  |  |  | - | 1.02 |
| XV |  |  |  |  |  | 1.07 |  |
| Mean | 1.38 | 1.20 | 1.27 | 1.16 | 1.15 | 1.27 | 1.09 |

Table 4. Contd.
| Species | *C. punctatus* | *A. insculpta* | *C. caeruleocinctus* | *C. londilensis* | *P. fuscus* |
| --- | --- | --- | --- | --- | --- |
| Stadia | L | L | L | L | L |
| II | 1.61 |  |  |  | 1.61 |
| III | 1.85 |  |  |  | 1.46 |
| IV | 1.28 | 1.32 |  | 1.40 | 1.42 |
| V | 1.57 | 1.74 | 1.23 | 1.27 | 1.31 |
| VI | 1.26 | 1.54 | 1.19 | 1.29 | 1.33 |
| VII | 1.35 | 1.52 | 1.27 | 1.26 | 1.26 |
| VIII | 1.32 | 1.46 | 1.23 | 1.31 | 1.25 |
| IX | 1.19 | 1.17 | 1.22 | 1.28 | 1.08 |
| X | 1.14 | 1.23 | 1.15 | 1.20 | 1.12 |
| XI | 1.06 |  | 1.13 | 1.10 | 1.11 |
| XII | 1.14 |  | 1.06 | 1.09 | 1.10 |
| XIII | 1.10 |  | 1.07 | 1.16 | 1.04 |
| XIV |  |  |  |  | 1.06 |
| XV |  |  |  |  | 1.08 |
| Mean | 1.32 | 1.43 | 1.17 | 1.24 | 1.23 |

The weight progression factor in nine species of millipedes including two polydesmid is presented in Table 5. In case of polydesmida the stadia increase in weight by a factor of three between two moults. Thus the weight does not double as would be expected according to the “Dyar-Hutchinson Rule” but in non-polydesmida the mean weight progression factor is 1.95 or 2 that follows the ‘Rule’. It is expected that addition of weight in successive stadia will be more when development time is short as in the case of polydesmida, and will be less in long development time found in non-polydesmida. Moreover, the weight progression factor in male and female is significantly different (P<0.05). This is due to accumulation of extra tissue (egg forming tissue) in female compared to male.

**Table 5.**
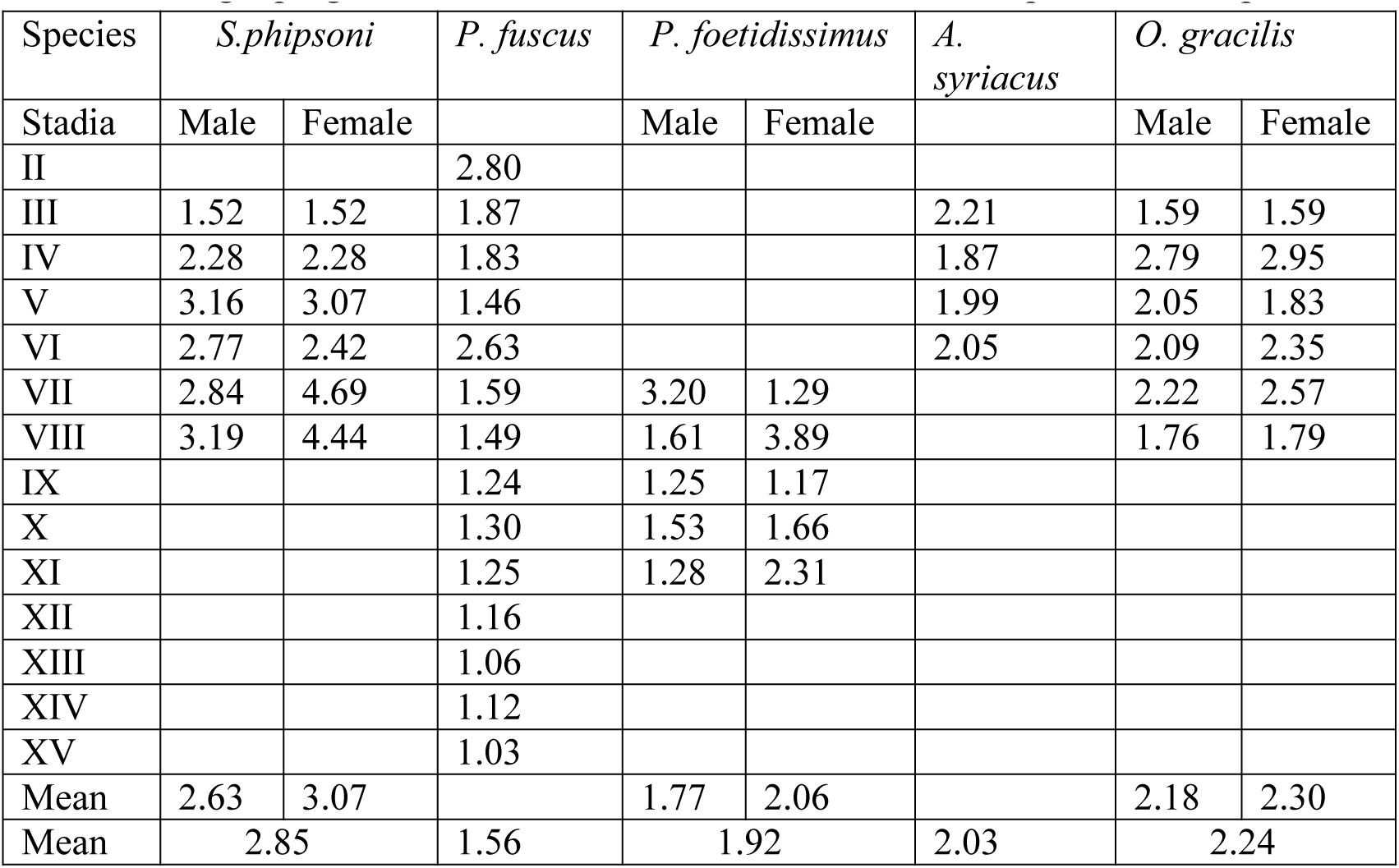
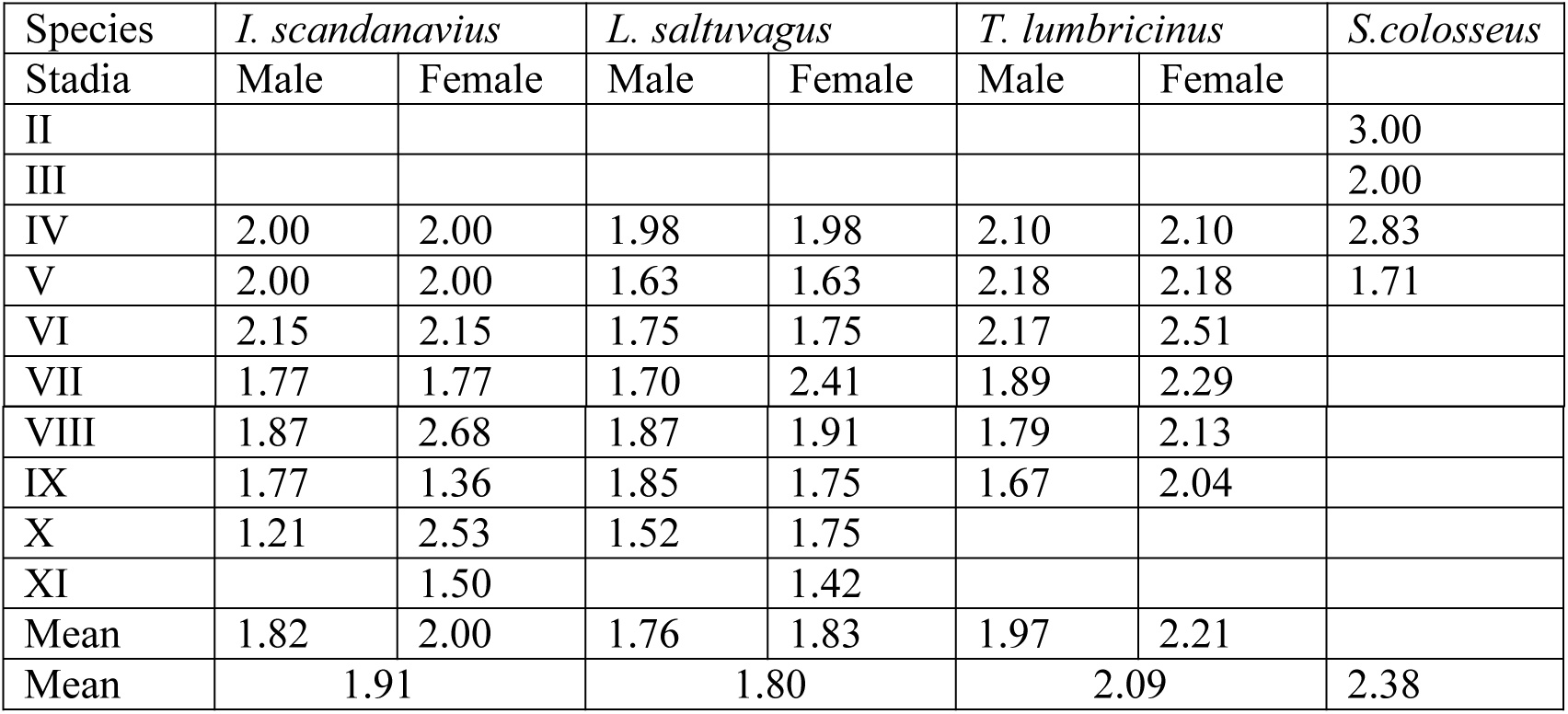
Weight progression factor of different stadia in different species of millipedes Species

## Discussion

There is no literature data on application of Dyar’s rule on millipede with which to compare my results. In the present study, there is close correspondence of the values of growth ratio of polydesmid millipede (mean 1.61, median 1.57) to Bodenheimer’s suggestion (1927) and that of non-polydesmid millipede (mean 1.23, median 1.23) to Przibram and Megusar rule (1.26). In most taxa, the modal values for overall size growth ranges from 1.15 to 1.50 such as around 1.50 for the larvae of holometabolous insects (Cole, 1980), 1.25 for hemimetabolous insects (Cole, 1980), 1.20 for decapods larvae (Rice, 1968), spiders (Enders, 1976) 1.16 for spiders (Humphreys, 1974) and trilobites (Fusco et al. 2012), 1.15 for lithobiomorpha centipedes (Albert, 1982) and 1.17 – 1.26 in Plecoptera (Elliott, 2009). The variations of Dyar’s value in the same animal group (here millipede – polydesmid and non-polydesmid) has also been observed by Cole (1980) in two insect group (holometabolous and hemimetabolous). Thus it can be concluded as recorded in other arthropod species, millipede growth in successive stadia is not in full agreement with Dyar’s rule as well as it varies from species to species or order to order. In this context, it can be mentioned that Dyar’s rule suggest that the growth increment should be constant for every instar but I found a systematic violation of the rule in that the growth progression varies from stadium to stadium. Moreover, there is a decreasing trend of Dyar’s value as the stadia advances (Albert, 1982 and the present study). Recently Hawes (2019) criticized Dyar’s law and detected several errors in the original data sheet of Dyar. Among 28 species of Lepidoptera, though number of instars varied from 4 to10 but Dyar’s values of those species not significantly varied. He proposed a new method of calculation of growth progression and applied this to calculate the growth progression value of all those 28 species. He elevated the value of growth progression of 28^th^ species, *Pyrrharctia isabella* having 10 instars to 2.3. This calculated values shows that Dyar’s value should be inversely proportional to the number of instars as found in several arthropods (Humphreys, 1974; Calvo and Molina, 2008; Elloitt, 2009; Nijhout and Callier, 2015; Grunert et al. 2015) including the present study. Dyar’s original data also support this hypothesis (Dyar’s value of butterflies having 5, 7 and 10 instar are 1.59, 1.42 and 1.30 respectively).

Several authors explain about the height of Dyar’s value. Przibram and Megusar (1912) commented that it equals to the cube root of 2 i. e. 1.26 and this is due to doubling of number of cells at each moult. But Bodenheimer (1927) claimed that growth progression factor may be higher that equals either1.26, 1.26^2^ = 1.6 or 1.26^3^ = 3. This seems to be true in case of millipede. But Horn and May (1977) and Maiorana (1978) commented that Dyar’s constant is of the same level as Hutchinson’s constant of about 1.28 which in congeneric sympatric species is the measure of difference in body length or length of feeding structure (Hutchinson, 1959).

Earlier Enders (1976), Cole (1980) and Hutchinson and Tongring (1984) advocated in favour of adaptive importance of Dyar’s rule. In this context, it is important to consider Dyar’s value from the functional point of view. Enders (1976) observed a correlation between the degree of locomotory activity and the height of Dyar’s constant. In different insect species it ranges from 1.16 to 1.69. In grazing herbivores which need not be very active show higher Dyar’s value but active predators have small Dyar’s value. This hypothesis is applicable in spiders also. Later Albert’s (1982) study on Lithobiidae is an affirmation of Ender’s hypothesis. The present study also supports Enders hypothesis because polydesmid are hygric in nature and confined to its resources for a long time with minimum movement (often form aggregation) show higher Dyar’s value. On the contrary, non-polydesmids are mesic-xeric adaptive species and more actively move for food resources and mate, show lower Dyar’s value.

Deviation from Dyar’s constant is observed in several species. This deviation can be calculated by Crosby’s growth rule which states that a difference exceeding ±10% in successive growth ratios indicate significant deviation.

Crosby’s growth ratio C_i_ = 100(r_i_ – r_(i-1)_) / r_(i-1)_, where, r = Dyar’s constant [Crossby, 1974a; Craig, 1975].

When, C_i_<10, there might be too many instars; when C_i_>10, some instars might be missing. In the second case (C_i_>10), it is expected that number of instars will be less with faster developmental time. In millipede this holds true. In polydesmid with C_i_>10 (in *S. phipsoni* C_i_ = 21.83) have only seven stadia with development time less than one year (Bhakat, 1989; Bhakat et al.1989; Banerjee, 1973) but in other non-polydesmid millipede with C_i_<10 (in *G. malayus* C_i_ = 3.02) have more than seven stadia and long development time (2-11 years) (Striganova and Mazantseva, 1979, Tracz, 1984, Bercovitz and Warburg, 1985). This variation of development time is related to an adaptation to habitat.

Some researchers reported Dyar’s rule for instar determination (Enrique, 2006, Klinberg and Zimmerman, 1992, Fink, 1984), though Gaines and Campbell (1935) did not recommend Dyar’s rule for instar determination as it may indicate false instars. I also think that number of instars (insect) or stadia (millipede) is species specific though different ecological and physiological factors may affect growth rate.

Several explanations have been put forwarded for specific values of Dyar’s constant in different species. Some suggest to ‘external cause’ by which species interact with its environment, such as competitive exclusion (Horn and May, 1977; Maiorana, 1978), food finding strategy (Enders, 1976), habitat stability (Cole, 1980), temperature and food (Klinberg and Zimmerman, 1992) and maximization of growth efficiency (Hutchinson and Tongring, 1984). In this case, a constant growth rate with a specific value is maintained by natural selection. On the other hand, some promoted to ‘internal cause’ which depend on species developmental system, anatomy and physiology, such as mechanism of intermoult hypodermal growth (Bennet-Clark, 1971; Freeman, 1991), physiology of moulting (Nijhout, 1981; Sehnal, 1985) and cell proliferation (Przibram and Megusar,1912; Bodenheimer, 1933). The two groups may be of different opinions but both the cause either external or internal have some weighed to ascertain the specific growth ratio or Dyar’s constant in a particular species or a species group as observed in millipedes.

The adaptive explanation of constancy of growth in successive instars by Hutchinson (1959) and later by Enders (1976) is that size increase (at least 28% or 1.26) has been selected to avoid competition between instars. Though the empirical evidence for Hutchinsons ratio between species of a guild has been challenged by Edie et al (1987) and MacNally (1988). Later ‘Investment Principle’ of Hutchinson et al. (1997) also provides adaptive explanation for Dyar’s rule providing the number of instars is optimized. They also explained the most common pattern of divergence from the rule. In the present study polydesmid millipede may be adapted to exploit ephemeral habitats (mostly available in the rainfall period) which appears patchily for a short period of time. To exploit such habitats millipede must grow rapidly with less number of stadia. Obviously rapid growth results higher Dyar’s value (>1.5). On the other hand, non-polydesmid millipedes exploit habitat that are available for a long period (comparatively in less hydrothermic condition) and may not require rapid development with more number of stadia. This results low growth ratio (<1.5).

When mean weight progression factor in between two groups (polydesmid and non-polydesmid) were compared, it is observed that in polydesmida, the weight progression factor is higher (2.55) compared to non-polydesmida (mean 1.97). In polydesmida, development time is short (within a year, normally 3-7 months) compared to non-polydesmid millipede (2-11 years). Duplication of mass with stadium in Julidae (non-polydesmida) is suggested by Blower and Gabbut (1964) and Blower (1970). So the weight progression factor in non-polydesmid millipede approves ‘Dyar-Hutchinson rule’ i. e. double their weight in successive moult. Polydesmid millipedes feed and grow only during relatively short periods (periods of rainfall) provided hydrothermic conditions are favourable. So higher weight progression factor in this group can be regarded as an adaptation of life under favourable conditions. On the contrary, as non-polydesmid millipedes are mesic-xeric adapted and can feed and grow beyond rainy season show lower weight progression factor. Andersson (1976, 1978b and 1980) and Albert (1982) observed that in Lithobiidae, the weight progression factor is lower when the species have a long development time. Humphreys (1974) also reported mean weight progression factor for wolf spider 1.57 having development time 548 days (consists of 16 size classes). Their observation is in concurrence with the present findings on millipedes. It is expected that weight growth of each stadium should be maximum to reach maturity where development time is short and vice versa. The above mentioned arthropods follow this rule. In millipedes, weight progression factor in females is significantly higher than that of male (t = 3.59, P<0.05). In wolf spider, weight progression factor in female is also higher compared to that of male (1.64 vs. 1.48) (Humphreys, 1974). It is due to more accumulation of reproductive tissue in female (mainly egg forming tissue) as the individuals are heading towards maturity.

These two groups of millipede i.e. polydesmid and non-polydesmid can be separated on the basis of several characters. Rapid development, early reproduction, small body size, semelparity, less tolerant to desiccation, hygric in habitat, fixed number of moults, strictly anamorphic development, all suggest polydesmid to be a r-adapted, time budget species while non-polydesmid millipede may be more k-adapted, energy budget species which is characterized by slow development, late reproduction, comparatively large body size, iteroparity, more tolerant to desiccation, mesic-xeric habitat, variable number of moults and hemianamorphic development (Blower, 1970; Bhakat, 2014). Present study added another character (Dyar’s constant value) to differentiate these two groups distinctly. In future, Dyar’ constant can be tested to differentiate organism belongs to r-selected and k-selected species.

## Acknowledgement

I am deeply indebted to my son Dr. Soumendranath Bhakat, Lund University, Sweden for constant support and my colleagues of Rampurhat College for their inspiration.

